# Isolation of *Actinomyces cricetomyis* sp. nov from orocervicofacial abscesses of African giant pouched rats (*Cricetomys ansorgei*)

**DOI:** 10.1101/2023.04.04.535624

**Authors:** Jordan D. Zehr, Rebecca J. Franklin-Guild, Scott V. Nguyen, Joseph R. Petrone, Rachael N. Labitt, Holly McQueary, Sebastian Llanos-Soto, Patrick K. Mitchell, Rebecca L. Tallmadge, Mason Jager, Timothy Wu, Alexander G. Ophir, Jonathan L. Jacobs, Anil Thachil, Bhupinder Singh, Laura B. Goodman

## Abstract

African giant pouched rats are notable both for their exceptional sense of smell and for how readily they can be trained, a combination that suits them to applications ranging from explosives detection to disease screening. A colony housed at a university animal care facility developed abscesses associated with the jaw and eye in multiple animals. The predominant bacterial species in each case was a catalase-positive *Actinomyces-*like Gram-positive bacillus. The isolates from different sites and animals matched each other genetically but had sequences and biochemical profiles inconsistent with previously described species of this group. The strains clustered together with 100% bootstrap support based on both 16S rRNA and whole genome alignment. Based on genomic metrics, the average nucleotide identity of the most closely related species is *Actinomyces respiraculi* ZJ750^T^ with 90.5% average nucleotide identity. Distinguishing biochemical characteristics include lack of nitrate reduction and hydrolysis of aesculin. Based on 16S rRNA, phylogenomic metric, and biochemical characterization, a novel species of the genus *Actinomyces* is proposed, namely *Actinomyces cricetomyis* sp. nov. The type strain is 186855^T^ (ATCC TSD-310^T^=LMG 32803^T^).

## INTRODUCTION

African giant pouched rats (*Cricetomys* spp.) are large terrestrial rodents from the family Nesomyidae, native to savannahs and evergreen forests of sub-Saharan Africa. These rodents stand out among other rodents for their relatively large olfactory cortex, bulbs, and ample olfactory receptor repertoire, which provide them with strong olfactory capabilities [1–4]. Due to their acute sense of smell, they have been successfully trained to diagnose tuberculosis cases in humans [5, 6], localize people trapped in collapsed structures [7], and employed in life-saving operations to detect anti-personnel landmines [8]. Additionally, they have showed potential for the detection of *Salmonella* in dried fecal samples [7]. Giant pouched rats have been identified as reservoirs of zoonotic bartonellae and hosts to trypanosomes [9, 10], as well as a potential source of Mpox transmission to humans [11–13]. However, only a few cases of diseases due to infectious agents have been described for this rodent, with reports of cestode cysts in wild-caught individuals [14] and induced illness due to Mpox virus in experimental settings [15]. While some bacterial-associated disease pathology in giant pouched rats has been reported, those of orocervicofacial origin have not been described.

The *Actinomyces* genus belongs to the family *Actinomycetaceae* that includes facultative and obligate anaerobic Gram-positive bacilli often present in the mucosae of animals and humans, such as the oral cavity, intestinal tract, and urogenital tract [16]. *Actinomyces* species are non-spore forming, rod-shaped, non-acid-fast bacteria with the ability to form filamentous branches [16, 17]. These bacteria are considered to be harmless organisms in humans under normal circumstances but are opportunistic pathogens when the integrity of the mucosa is breached through trauma, surgery or infection [18]. Under these conditions, *Actinomyces* can invade local tissue and adjacent organs causing actinomycosis, an infrequent, non-contagious, chronic pathology that involves the continuous and progressive formation of abscesses [19]. Actinomycosis has different presentations depending on the anatomical sites affected, namely orocervicofacial, thoracic, abdominopelvic, central nervous system, musculoskeletal, and disseminated [20]. Although prognosis is very favorable for most presentations of the disease following prolonged antibiotic treatment, diagnosis remains challenging due to its unspecific symptoms, wide range of different clinical presentations, and strong similarity with pathologies such as tuberculosis and malignancy [21].

## METHODS

### Isolation and ecology

Southern giant pouched rats (*Cricetomys ansorgei*) 1 and 2 were captured from a wild population at a field site in Morogoro, Tanzania (6.8304° S, 37.6706° E) and were founding members of the colony at Cornell University in Ithaca, NY USA. Animals were captured using collapsible medium sized Havahart traps (Model 1090). Pouched rat 3 was born in captivity at Cornell University. Animals were housed singly in standard rabbit cages (61 x 61 x 41 cm) in a same-sex room in an accredited animal facility. All work and maintenance of the animal colony followed American Society of Mammalogists guidelines [22] and was approved under US Army Medical Research and Materiel Command (USAMRMC) Animal Care and Use Review Office (ACURO) and the Cornell University Institutional Animal Care and Use Committee. The first animal developed otitis media in Oct. 2017. There were three isolates recovered from the second animal, one in 2018 (orbital abscess, resulting in eye removal), one in 2019 (orbital abscess, resulting in second eye removal), and one in 2020 (orbital abscess post-eye removal). The third animal developed a maxillary abscess in July 2021. In all cases, swabs were submitted for culture and sensitivity as part of standard of care treatment. Gram staining of abscess material from all three individuals showed moderate to high presence of Gram-positive thin branching rods. The initial aerobic bacterial cultures on these specimens were performed using standard procedures of the Cornell University Animal Health Diagnostic Center (AHDC) Bacteriology lab. Specimens were plated to Trypticase soy agar with 5% sheep blood, Levine eosin methylene blue (EMB) agar and Columbia cephalothin nalidixic acid (CNA) agar and struck for isolation. Culture plates were incubated overnight at 33-37°C in 6% ± 1 CO_2_ and evaluated at 18-24h and again at two days following plating. Culture colonies deemed potentially significant were identified by MALDI-TOF mass spectrometry using the Bruker MALDI Biotyper ® System (Bruker Scientific), along with conventional biochemical screens and 16S rRNA sequencing for these isolates. Archival of the Actinomyces-like organisms was performed by growing them in trypticase soy broth overnight at 33-37°C in 6% ± 1 CO_2_, mixing the broth in equal amounts with 30% glycerol in a labeled cryovial and storing at -80 ± 3°C. All subsequent work in this study was completed with isolates regrown from the archived organisms

### Isolate genome sequencing and assembly QC

Genomic DNA was extracted from an overnight culture of each strain using the Qiagen DNA Blood and Tissue extraction kit following the manufacturer’s protocol. The DNA from isolates of the first two animals was then prepared for whole genome sequencing using the DNA Prep library kit from Illumina, and sequencing was performed on the Illumina MiSeq with 2x250 bp chemistry. Raw FastQ files were uploaded to NCBI BioProject PRJNA775986 (see Table 1 for individual strain accessions). Quality control using FastQC version 0.11.8 [23] was performed on the reads, followed by trimming to remove sequencing adapters and low-quality bases using trimmomatic version 0.39 [24] (parameters: LEADING:3 TRAILING:3 SLIDINGWINDOW:4:15 MINLEN:200 –phred33). Reads were then *de novo* assembled using SPAdes version 3.15.2 [25], and assemblies were checked for quality controls using QUAST version 5.1.0rc1 [26].

**Table 1.**

| Strain | Organism | Isolation Host | Isolation Source | Collection Date | Accession |
| --- | --- | --- | --- | --- | --- |
| 186855<sup>T</sup> | *Actinomyces cricetomyis* sp. nov | *Cricetomys ansorgei* | Ear | October 2017 | GCF_060694225.1 |
| 187325 | *Actinomyces cricetomyis* sp. nov | *Cricetomys ansorgei* | Orbital abscess prior to eye removal | December 2018 | GCF_023667625.1 |
| 217892 | *Actinomyces cricetomyis* sp. nov | *Cricetomys ansorgei* | orbital abscess post-eye removal | October 2019 | GCF_023667585.1 |
| AC-20-1 | *Actinomyces cricetomyis* sp. nov | *Cricetomys ansorgei* | orbital abscess post-eye removal | June 2020 | GCF_023369555.1 |
| W5033 | *Actinomyces* sp. | *Homo sapiens* | nasopharyngeal swab | 2025 | GCF_051303165.1 |
| ZJ750 | *Actinomyces respiraculi* | *Marmota himalayana* | respiratory tract | July 2018 | GCF_014595995.2 |
| ZJ751 | *Actinomyces respiraculi* | *Marmota himalayana* | NA | July 2018 | GCF_011751965.1 |
| CSUR P35 | *Actinomyces timonensis* | *Homo sapiens* | clinical osteo-articular sample | 2007 | GCF_000295095.1 |
| DSM 20671 | *Actinomyces denticolens* | *Bos taurus* | dental plaque | NA | GCF_001995055.1 |
| CGMCC 4.7372 | *Actinomyces gaoshouyii* | *Ochotona curzoniae* | intestinal contents | NA | GCF_014646495.1 |
| zg-325 | *Actinomyces marmotae* | *Marmota himalayana* | respiratory tract | August 2018 | GCF_013177295.1 |
| PK606 | *Actinomyces stomatis* | *Homo sapiens* | oral cavity | NA | GCF_031292835.1 |
| VPI D163E-3 | *Actinomyces acetigenes* | *Homo sapiens* | gingival crevice | NA | GCF_031296635.1 |
| CCUG 34288 | *Actinomyces oris* | *Homo sapiens* | sputum | NA | GCF_006546825.1 |
| zg-993 | *Actinomyces trachealis* | *Marmota himalayana* | respiratory tract | July 2018 | GCF_015711475.1 |
| CCUG 61299 | *Actinomyces weissii* | *Canis familiaris* | oral cavity | NA | GCF_016598775.1 |
| JCM 18848 | *Actinomyces haliotis* | *Haliotis discus* subsp. *hannai* | gut | January 2013 | GCF_015355765.1 |
| CCUG 36733 | *Actinomyces radidentis* | *Homo sapiens* | infected root canal | 1966 | GCF_001553565.1 |
| ZJ34 | *Actinomyces faecalis* | *Marmota himalayana* | feces | July 2018 | GCF_013184985.2 |
| DSM 15434 | *Actinomyces urogenitalis* | *Homo sapiens* | vaginal secretions | NA | GCF_000159035.1 |
| DSM 19115 | *Actinomyces dentalis* | *Homo sapiens* | dental abscess | 2003 | GCF_000429225.1 |
| DSM 15435 | *Actinomyces bowdenii* | *Canis familiaris* | mandible abscess | NA | GCF_017592395.1 |
| ATCC 49928 | *Actinomyces slackii* | *Bos taurus* | dental plaque | NA | GCF_000428685.1 |

*Actinomyces* NCBI genome accessions used in this study along with isolation metadata.

In a separate experiment, genomic DNA from strain 186855^T^ (=ATCC TSD-310^T^) was extracted as previously described [27]. For the short-read Illumina dataset, a paired-end sequencing library for strain 186855^T^ was sequenced on an Illumina NextSeq 2000 using the P2, 300-cycle (2x150bp) chemistry. Genomic DNA from the same extraction was barcoded using an EXP-NBD196 kit and sequenced on an Oxford Nanopore Technology (ONT) GridION sequencer with R9.4.1 chemistry. ONT reads were base called using the high-accuracy settings, demultiplexed and barcode-trimmed. A hybrid assembly was produced using Trycycler v0.5.4 [28] and polished by Medaka v1.11.3, Polypolish v0.6.0 [29], and pypolca v0.3.1 [30]. A combination of Flye v2.9.3-b1797 [31], miniasm v0.3-r179, and raven v1.8.3 were used to generate the input assemblies for Trycycler. This hybrid approach resulted in a closed, finished genome composed of a single replicon at a length of 2,707,859 nucleotides (72.64% GC content), from NCBI (BioProject: PRJNA1481675) and the ATCC Genome Portal (Shttps://genomes.atcc.org/genomes/c202f05248c5406e) [32] GCF_060694225.1.

### 16S Phylogeny and genomic SNP tree

RNAmmer version 1.2 [33] was used to identify complete length 16S rRNA gene sequences (1551bp) from assemblies of animals 1 and 2, and bedtools [34] getfasta version 2.29.2 was used to extract the 16S rRNA gene sequences. NCBI BLAST was used to pull down the top 100 homologous sequences, with 186855^T^ as the query. MAFFT version v7.526 [35] was then used to align the 16S rRNA gene sequences, and RAxML-ng version 1.2.2 [36] was used to build the 16S phylogenetic tree using the GTR+G algorithm (parameters --all --model GTR+G --bs-trees 1000 --seed 1234 --threads 8).

A k-mer-based phylogenetic analysis was performed using CLC Genomics Workbench v26 (Qiagen, Aarhus, Denmark). Genome assemblies of the proposed type strain, additional isolates collected from this study, and closely related *Actinomyces* type strains as identified through the Type (Strain) Genome Server (TYGS) [37] were analyzed by k-mer analysis. Pairwise genomic distances were calculated using the Feature Frequency Profile (FFP) method with a k-mer size of 16 and considering both DNA strands. This alignment-free approach was used to construct a phylogenetic tree depicting relationships among the analyzed genomes.

### Whole-Genome comparisons

The Type (Strain) Genome Server [37] was implemented to calculate digital DNA-DNA hybridization (dDDH) values of strain 186855^T^. Related type strains identified by TYGS were then used in an all vs all average nucleotide identity (ANI) comparison as implemented in ANIclustermap (https://github.com/moshi4/ANIclustermap). The accession for strains used in this study are listed in Table 1.

### Morphological, physiochemical and biochemical characterization

All isolates were analyzed by MALDI-TOF and 16S rRNA sequencing at the AHDC. Biochemical tests of the four novel strains and two close relatives were conducted using API ZYM system and Rapid ID 32A according to manufacturer’s instructions. Both the manually inoculated API ZYM and Rapid ID 32A strips were incubated at 33-37°C in aerobic conditions for 4 to 4.5 hours prior to being read visually, following the addition of necessary reagents. All reactions were read in duplicate by two separate technicians and recorded as the consensus of each result. Additional culture media and temperature testing was performed to determine optimal growth conditions, as well as conventional biochemical screening, following the laboratory’s standard operating procedures.

## RESULTS AND DISCUSSION

### Pathological changes associated with infection

Strain 186855^T^ (Table 1) was the predominant bacterial strain isolated from purulent exudate draining from the ear canal of a male pouched rat submitted for routine clinical culture. The exudate originated from an abscess of the jaw with associated soft tissue swelling, extensive osteomyelitis, and pathologic fracture. Histopathology showed intralesional Gram-positive filamentous rods with Splendore-Hoeppli material, consistent with actinomycosis in other species [38], and susceptibility testing showed that the isolate was resistant to enrofloxacin and penicillin. Strain 187325 was isolated from a different male pouched rat housed in the same room, which developed surgical site abscesses at three, fifteen, and twenty-three months after surgical removal of the eye due to a corneal ulcer that was refractory to treatment. Histopathology of the removed eye revealed long filamentous rods present within the cornea. Initial attempts for identification revealed white non-hemolytic colonies at culture under aerobic conditions. Strain 154992 was the predominant aerobic bacterial strain isolated from a third male pouched rat housed in the same room. This animal exhibited an acute onset of inappetence, tremors and difficulty walking. Despite treatment, the animal continued to decline and was euthanized. On necropsy, a purulent material was expressed from the left caudal maxilla which was submitted for aerobic and anaerobic culture. The predominant organism in aerobic culture was a white, non-hemolytic colony resembling those in the previous cases, and histopathology results were analogous to those on the animal where strain 186855^T^ was recovered.

### Biochemistry

The four novel strains proposed herein could be differentiated from *A. radicidentisi* CCUG 36733 and *A. urogenitalis* CCUG 38702^T^ comparison strains with the valine arylamidase, β-glucuronidase, and N-acetyl β-glucosaminidase biochemical tests values (Table 2 and Table 3). Additional biochemical features were used in comparison to data from Zhou *et al* [39]. Yet, some degree of phenotypic heterogeneity is observed among the proposed novel strains in Table 2, particularly for histidine arylamidase activity. Such variation in biochemical characteristics has previously been reported among catalase-positive *Actinomyces* strains. Notably, all four novel strains were negative for tyrosine arylamidase and weakly positive for β-glucuronidase when compared to other closely related *Actinomyces* species. The novel strains were also positive for α-arabinosidase, arginine arylamidase, leucyl glycine arylamidase, and serine arylamidase in comparison to the closest identified species, *Actinomyces respiraculi* [40].

**Table 2.**

| Characteristic | 1 | 2 | 3 | 4 | 5 | 6 | 7 | 8 | 9 | 10 | 11 | 12 | 13 |
| --- | --- | --- | --- | --- | --- | --- | --- | --- | --- | --- | --- | --- | --- |
|  | Enzyme activity (API ZYM): |  |  |  |  |  |  |  |  |  |  |  |  |
| Alkaline Phosphatase | - | - | - | - | - | - | - | - | - | - | - | - | - |
| Esterase (C 4) | - | - | - | w | w | - | - | - | - | - | + | - | - |
| Esterase Lipase (C 8) | - | - | - | - | - | - | - | - | - | - | + | - | w |
| Lipase (C 14) | - | - | - | - | - | - | - | - | - | - | - | - | - |
| Leucine arylamidase | + | + | + | + | + | + | + | + | + | + | + | + | + |
| Valine arylamidase | - | - | - | - | - | - | + | - | w | w | + | w | w |
| Cystine arylamidase | - | - | - | - | - | - | - | - | w | - | - | - | w |
| Trypsin | - | - | - | - | - | - | - | - | - | - | - | - | - |
| α-chymotrypsin | - | - | - | - | - | - | - | - | - | - | - | - | - |
| Acid phosphatase | - | - | - | - | - | - | - | - | - | - | - | - | - |
| Naphthol-AS-BI-phosphohydrolase | - | - | - | - | - | - | - | - | - | - | - | - | - |
| α-galactosidase | + | + | + | + | + | - | - | + | - | + | + | + | - |
| β-galactosidase | + | + | + | + | + | - | + | + | + | + | + | + | + |
| β-glucuronidase | - | w | w | w | w | - | - | - | - | - | - | - | - |
| α-glucosidase | + | + | + | + | + | + | - | + | + | + | + | + | - |
| β-glucosidase | + | + | + | + | + | + | + | + | + | + | + | + | + |
| N-acetyl β-glucosaminidase | + | + | + | + | + | + | + | - | + | + | + | + | + |
| α-mannosidase | - | - | - | - | - | - | - | - | - | w | + | - | - |
| α-fucosidase | - | - | - | - | - | - | - | - | - | - | - | - | - |

Characteristics that differentiate *A. cricetomyis* strains 186855<sup>T</sup>, 217892, 187325, AC-20-1, and 154992 from the type strains of related species of *Actinomyces*. Biochemical features from Zhou *et al.* were also used in this analysis. Strains: 1, 186855<sup>T</sup>; 2, 217892; 3, 187325; 4, AC-20-1; 5, 154992; 6, *A. respiraculi* ZJ750<sup>T</sup>; 7, *A. trachealis* zg-993<sup>T</sup>; 8, *A. radidentis* CCUG 36733<sup>T</sup>; 9, *A. weissii* CCUG 61299<sup>T</sup>; 10, *A. urogenitalis* CCUG 38702<sup>T</sup>; 11, *A. faecalis* ZJ34<sup>T</sup>; 12, *A. bowdenii* DSM 15435<sup>T</sup>; 13, *A. bovis* CCTCC AB2010168<sup>T</sup>; -, Negative; +, positive; w, weakly positive.

**Table 3.**
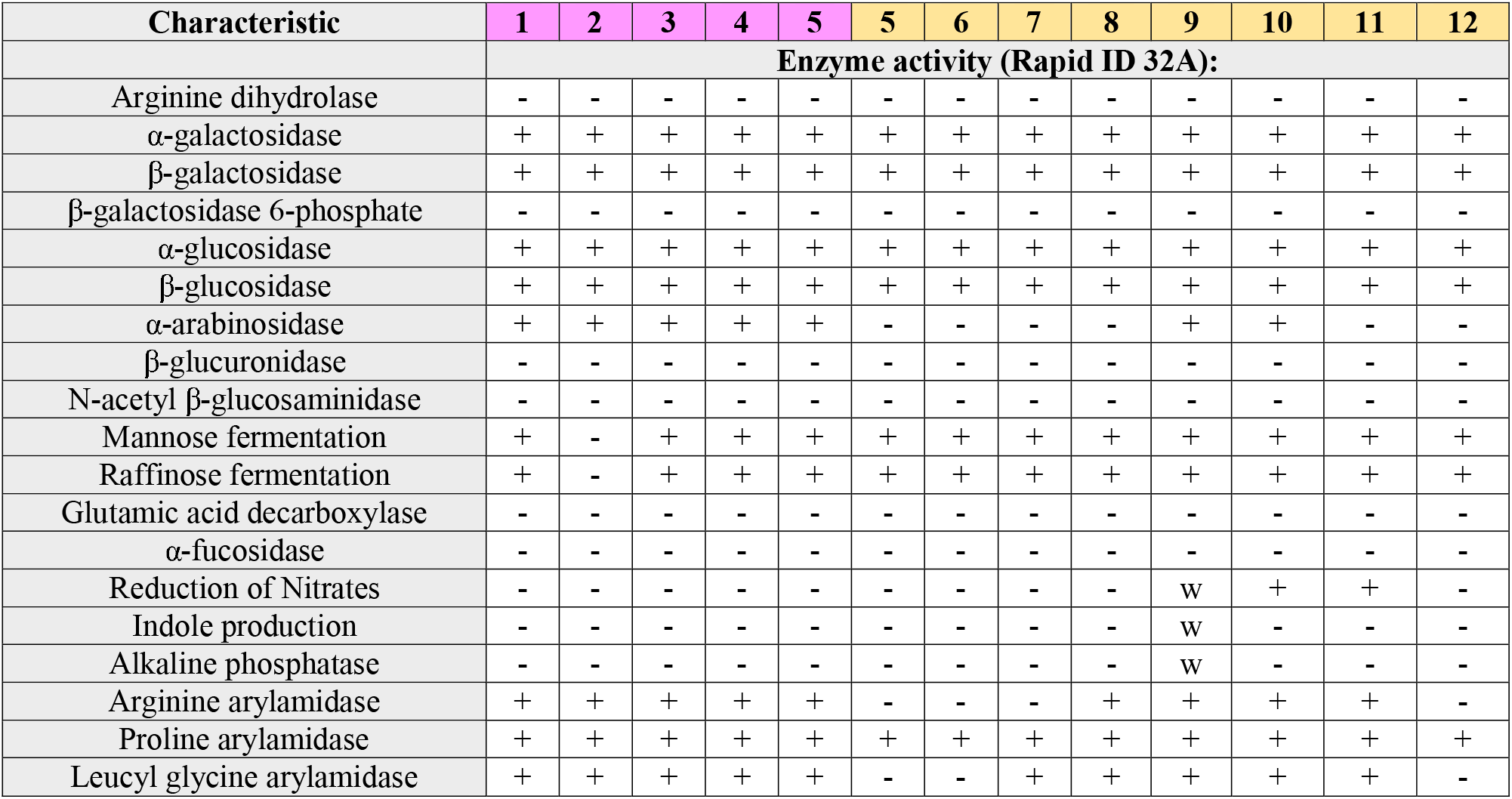
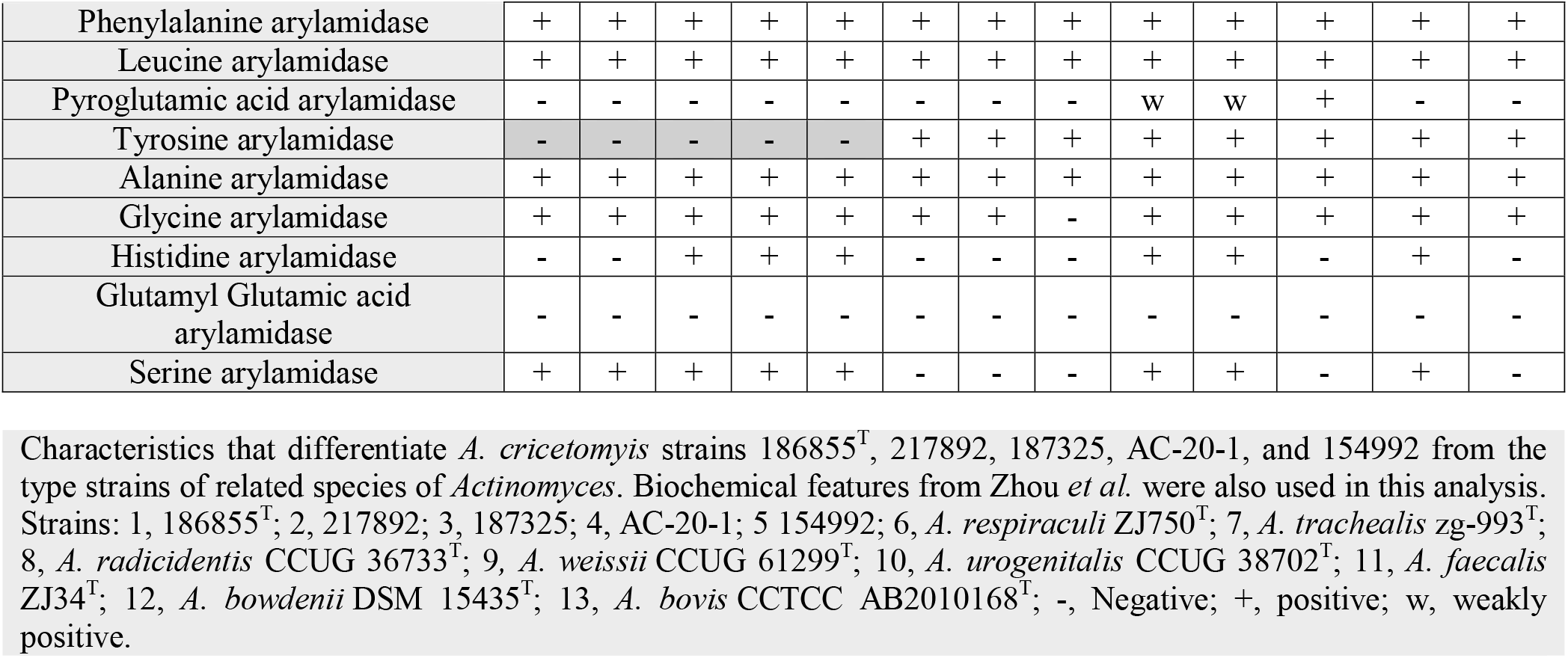

### 16S rRNA analysis

Initial analysis of partial 16S rRNA gene sequences determined through Sanger sequencing identified all the isolates as *Actinomyces sp*. with best matches being *Actinomyces bowdenii* DSM 15435^T^ (95-96% identity), *Actinomyces weissii* CCUG 61299^T^ (95% identity), and *Actinomyces bovis* DSM 43014^T^ (95% identity) based on BLAST. Phylogenetic analysis of the full-length 16S rRNA extracted from the shotgun sequence is shown in Fig. 1. The robust separation of the three *Actinomyces cricetomyis* strains from all other taxa, supported by 100% bootstrap replication, provides an initial line of evidence for their distinctiveness; however, the poor bootstrap support throughout the remainder of the tree is consistent with well-documented limitations of 16S rRNA as a phylogenetic marker for *Actinomyces* [41–43]. Three principal phylogenetic clusters can be discerned within the genus, but these are not strongly supported by bootstrap analysis, and inter-species 16S rRNA sequence similarities are often high enough to confound discrimination [44]. The 16S tree therefore serves primarily to confirm that the novel strains form a coherent, distinct group.

**Fig. 1.**
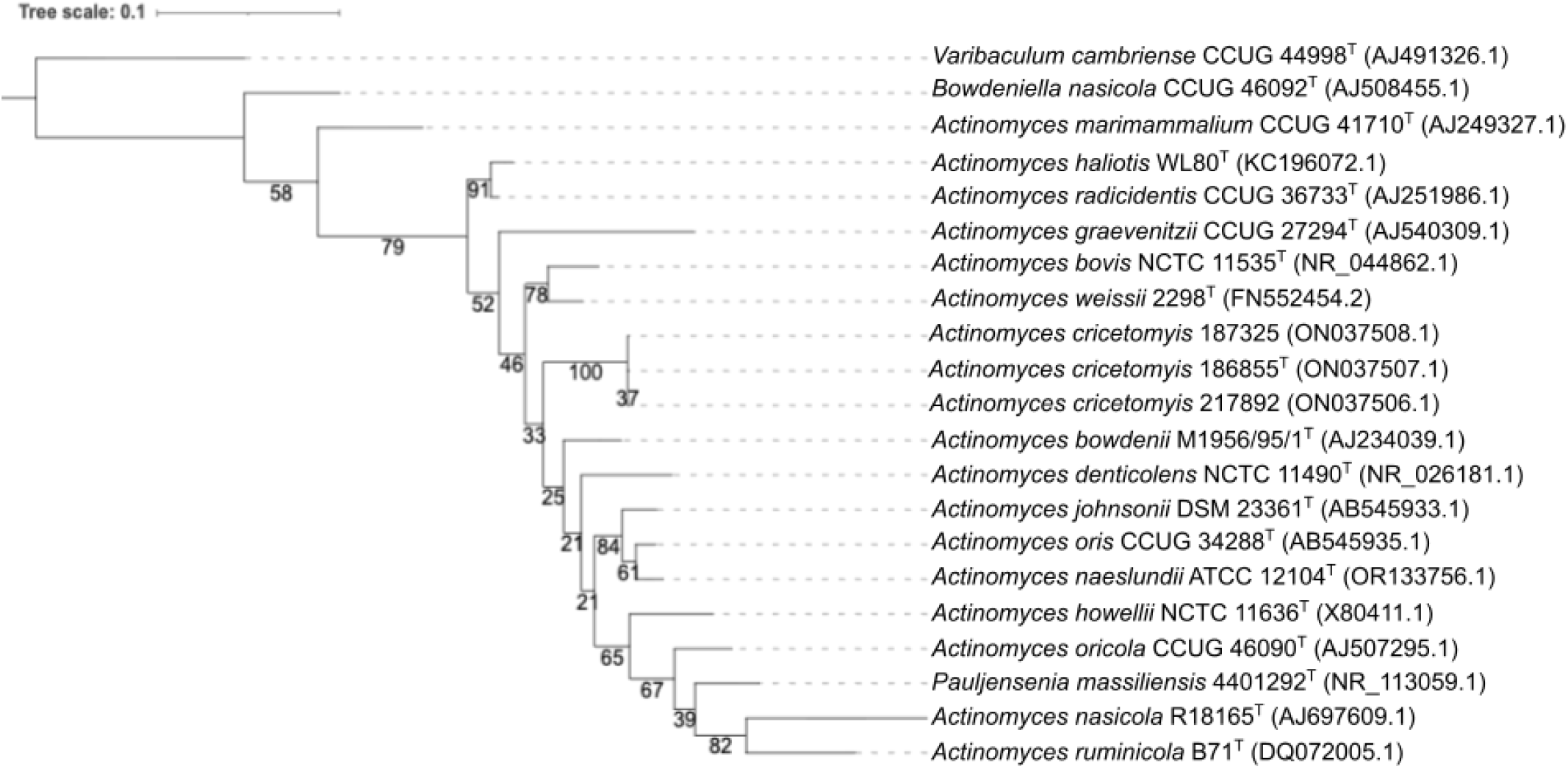
16S Phylogeny Phylogenetic tree of 16S rRNA gene from proposed *Actinomyces cricetomyis* (3 isolates; sp. 217892, sp. 186855T, and sp. 187325), along with diverse *Actinomyces* species. The tree was inferred with raxml-ng with 1000 bootstrap replicates and rooted with *Varibaculum cambriense* CCUG 44998^T^ (AJ491326.1). The scale bar indicates the branch lengths within the tree

### Genomic analysis

The average genome size of these strains are 2.7 Mb with GC content of 72.4%. The hybrid assembly of strain 186855^T^ (GCF_060694225.1) resulted in a single, circular genome at 2707859 bp with 2,316 predicted genes of which 2,232 genes are protein-coding. 51 tRNAs were identified, and 3 rRNA for each (5S, 16S, and 23S) clusters were annotated. An alignment-free k-mer tree inferred with *Actinomyces* genomes further demonstrates monophyly of the *Actinomyces cricetomyis* strains, distinct from the closest related species *A. respiraculi* (Fig. 2). The k-mer tree provides substantially greater resolution than the 16S rRNA tree, consistent with the broader finding that whole-genome approaches outperform single-marker analyses for delineating species within *Actinomycetaceae* [44, 45]. The *Actinomyces cricetomyis* strains form a strongly supported monophyletic group, a relationship also reflected in the ANI comparisons (Table 3), where *A. respiraculi* and related strains return the highest non-conspecific ANI values against our type strain. This congruence between the k-mer tree phylogeny and the pairwise genomic distance metrics provides mutual corroboration for the placement of these strains. Notably, the clustering of the isolates recovered from separate animals over a four-year period from distinct anatomical sites with pairwise ANI values >99.8% is consistent with a single persistent lineage in this animal colony rather than repeated independent introductions, which may have implications for the clinical management of affected animals and infection control within captive colonies. The genome tree further illustrates the broad genomic diversity within *Actinomyces* as currently defined, consistent with longstanding recognition that the genus is polyphyletic and likely warrants further subdivision [44]. The placement of *A. cricetomyis* sp. nov. as a well-supported, deeply branching lineage distinct from all currently described species further argues for its recognition as a novel taxon.

**Fig. 2.**
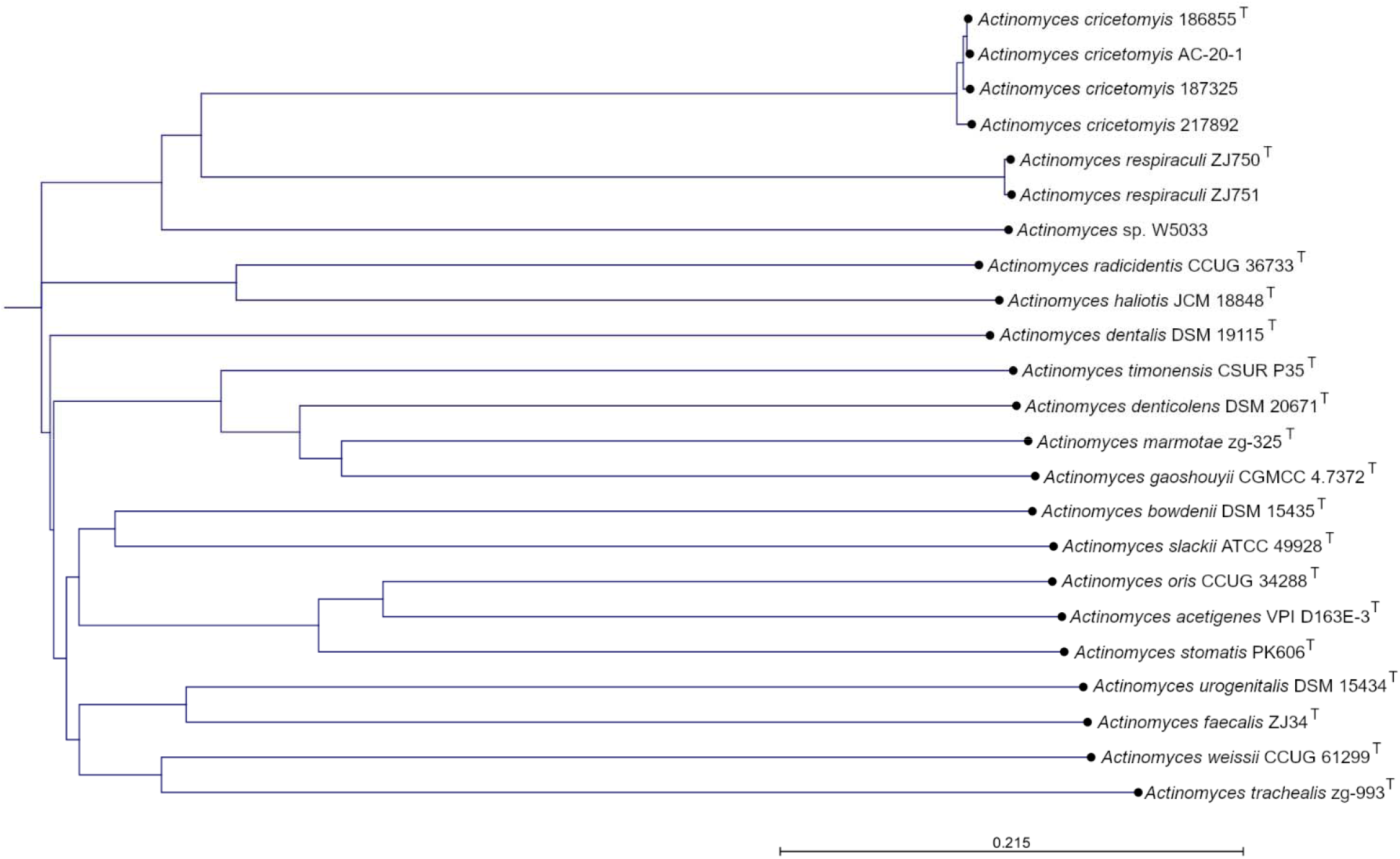
Genome phylogeny Phylogenetic tree of the proposed *Actinomyces cricetomyis* to other related *Actinomyces* species. The tree was constructed in CLC Genomics v24 using an alignment-free *k*-mer–based approach of all genomes. The scale bar indicates the branch lengths within the tree.

An all vs all ANI comparison of type strains identified by TYGS also display less than 95% ANI similarity of the proposed *A. cricetomyis* to related *Actinomyces* (Figure 3). Published metrics indicate ANI values < 95% suggest a species delineation [46], suggesting the genomes we report herein are a novel species compared to those publicly available. Recent informal genomic characterizations have described *Actinomyces acetigenes* and *Actinomyces stomatis* from the human oral cavity using ANI, dDDH, and pangenome approaches [47], and *Schaalia dentiphila* from the same ecological niche, with pangenome and genomic island analysis further illuminating functional diversity within this family [48]. These parallel efforts underscore the genomic heterogeneity and ongoing taxonomic revision within *Actinomycetaceae*.

**Fig. 3.**
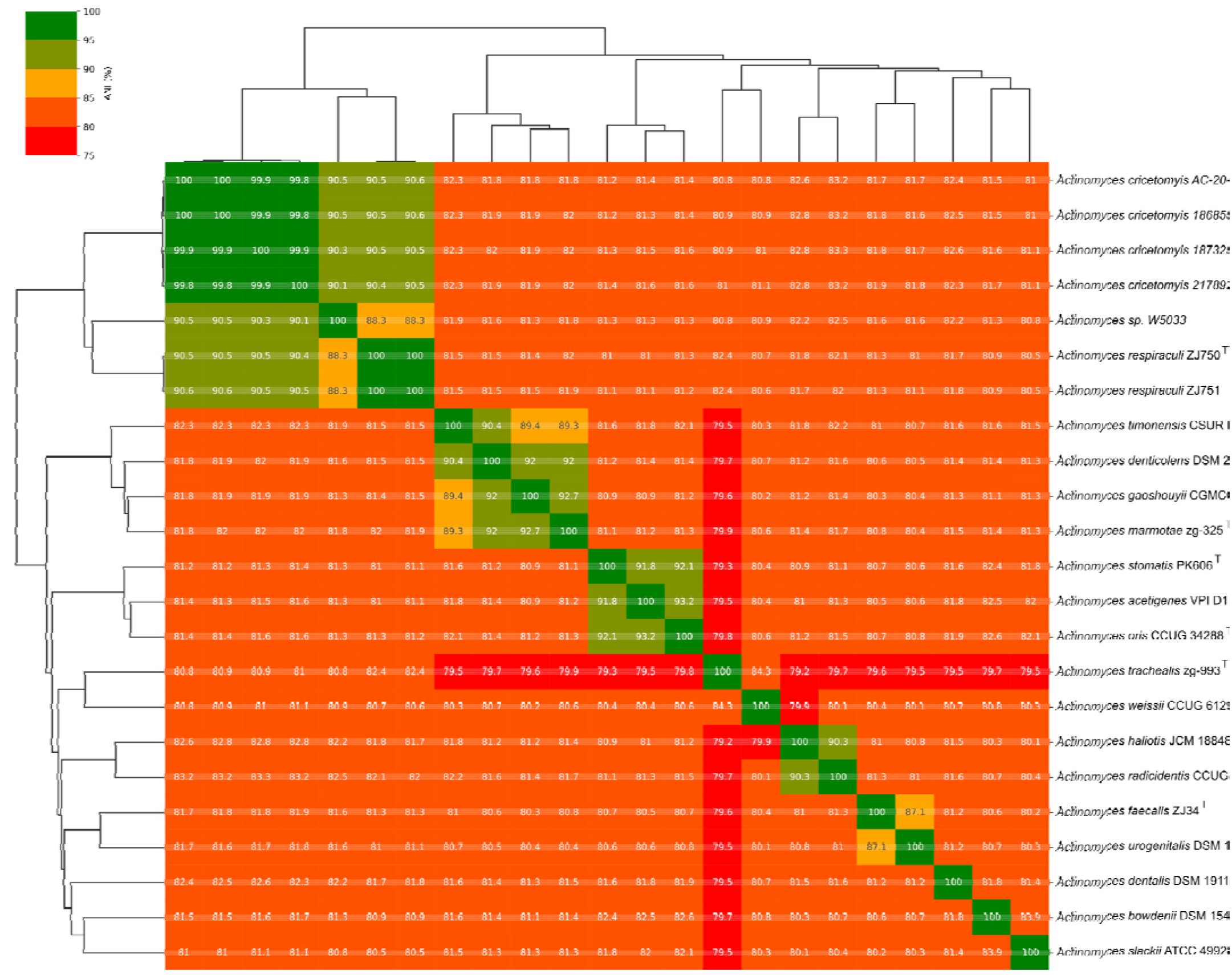
ANI Matrix FastANI visualization of the proposed *Actinomyces cricetomyis* strains to other closely related Actinomyces strains and related type strains of other *Actinomyces* species. All members of *A. cricetomyis* show less than 95% ANI to other species.

*Actinomyces respiraculi* ZJ750^T^ (GCF_014595995.2) was identified as the most closely related *Actinomyces* species to strain 186855^T^ with a dDDH of 37.8% when utilising formula d_4_ (Table 4). With a species delineation threshold published at ≤ 70% [46, 49], this suggests that the novel genomes reported herein are indeed a new species. It is curious that *Actinomyces respiraculi* ZJ750^T^ the most closely related species to the type strain reported herein according to ANI and dDDH, respectively, is associated with respiratory infections [50]. This is contrasted with the *Actinomyces* genomes isolated from the animals in this study were derived from the ear and eye abscesses of infected animals. Further, our findings suggest that while these *Actinomyces* species are the most closely related, they all have been isolated from different hosts (*Actinomyces respiraculi* [GCF_014595995.2] from a marmot and *Actinomyces cricetomyis* sp. nov [186855^T^] from pouched rats, suggesting a broad host range and infection tropism for *Actinomyces*. A closely related, currently undescribed *Actinomyces* strain, *Actinomyces* sp. W5033, was also identified by ANI analysis. According to available NCBI metadata, strain W5033 was isolated from a human nasopharyngeal swab. To our knowledge, no formal taxonomic description or published characterization of this strain is currently available, and its clinical significance remains unknown. Additionally, the other related *Actinomyces* strains also show a broad host range. See Table 1.

**Table 4.**
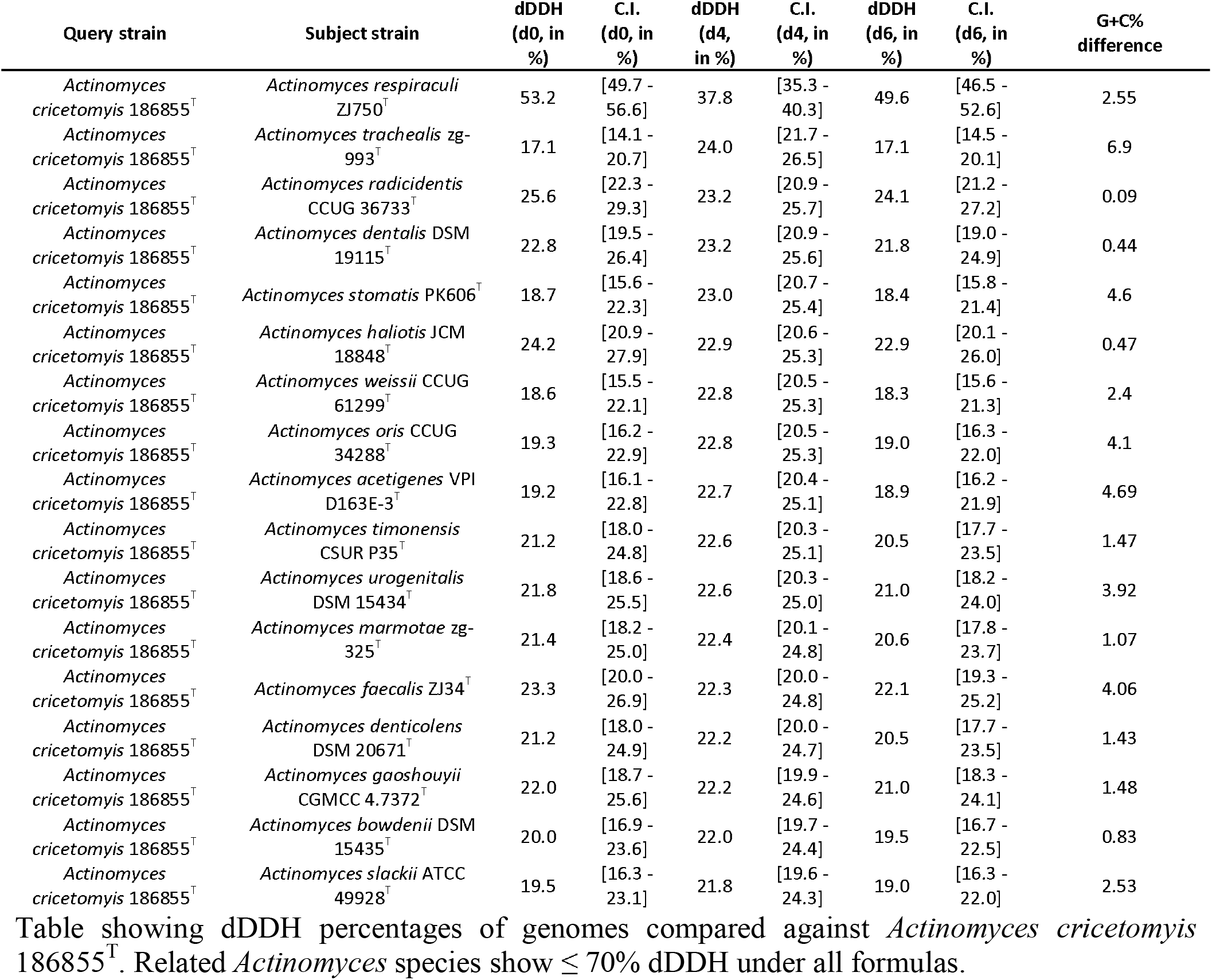

The combined polyphasic results of the phenotypic, phylogenetic, and genomic analyses support the conclusion that strain 186855^T^ represents as a member of a novel species within *Actinomyces*. Phylogenetic reconstruction based on 16S rRNA gene sequences and alignment-free k-mer analysis show that the isolated strains form a distinct lineage separate from other currently described *Actinomyces* species (Figure 1). Furthermore, genome relatedness indices, including ANI and dDDH, are below the accepted thresholds for species delineation with its closest phylogenetic neighbors. Phenotypic characterizations also identified biochemical features that are differentiated from strain 18655^T^ from related type strains and collectively, these data support the recognition that these strains represent a novel species within *Actinomyces* for which the name *Actinomyces cricetomyis* sp. nov. is proposed. Strain 186855^T^ is designated as the type strain for *Actinomyces cricetomyis* sp. nov.

### Global ecological distribution

The ecological distribution patterns of 186855^T^ were addressed using Sandpiper [51] (v2.1.1) (https://sandpiper.qut.edu.au/), an interface that uses the SingleM tool to accurately map metagenomic reads to genomes. At the time of the analysis, 913,071 distinct metagenomes (from 42, 716 projects) were screened and made available through Sandpiper, with taxonomy annotations matching the Genome Taxonmy Database (GTDB) [52] release 232. The whole genome sequence GCF_060694225.1 was used as query, and results were visualized on the Sandpiper website to assess the global distribution patterns (occurrence and relative abundance) and habitat preferences of 186855^T^ (https://sandpiper.qut.edu.au/taxonomy/s Actinomyces%20sp023369555). Of the 37 runs with matching samples, 9 runs had ≥ 0.23% relative abundance, with 9 runs reporting geographic location associations. The top run (ERR6178868) had a relative abundance of 2.77%, from the oral environment of dental calculus from four Chalcolithic individuals from the Iberian peninsula (ca. 4500-5000 BP) [53]. The second most abundant run came from a study of the human oral microbiomes from hunter-gatherer and subsistence farmers compared to highly industrialized populations [54]. Hits from 186855^T^ were also recovered from chicken (https://www.ncbi.nlm.nih.gov/bioproject/PRJNA1039992) and sheep gut environments [55], and sewage from Kolkata (https://www.ncbi.nlm.nih.gov/bioproject/PRJDB15925).

## Abbreviations

ANI: average nucleotide identity
dDDH: digital DNA-DNA hybridization

## Data Availability

The Genbank accession numbers for the 16S rRNA gene sequences of strains 186855^T^, 187325, 217892, and 154992 are ON037507.1, ON037508.1, ON037506.1, and OQ458734, respectively. The Genbank accession numbers for the genome sequences for strains 186855^T^, 187325, 217892, and AC-20-1 are GCF_060694225.1, GCF_023667625.1, GCF_023369555.1, and GCF_023667585.1, respectively. The BioSample Accession numbers for strains 186855^T^, 187325, 217892, and AC-20-1 are SAMN61087245, SAMN22657059, SAMN22657060, and SAMN15639644, respectively. The SRA accession numbers for strains 186855^T^, 187325, 217892, and AC-20-1 are SRR17017921, SRR17017920, SRR17017919, and SRR12413902 respectively.

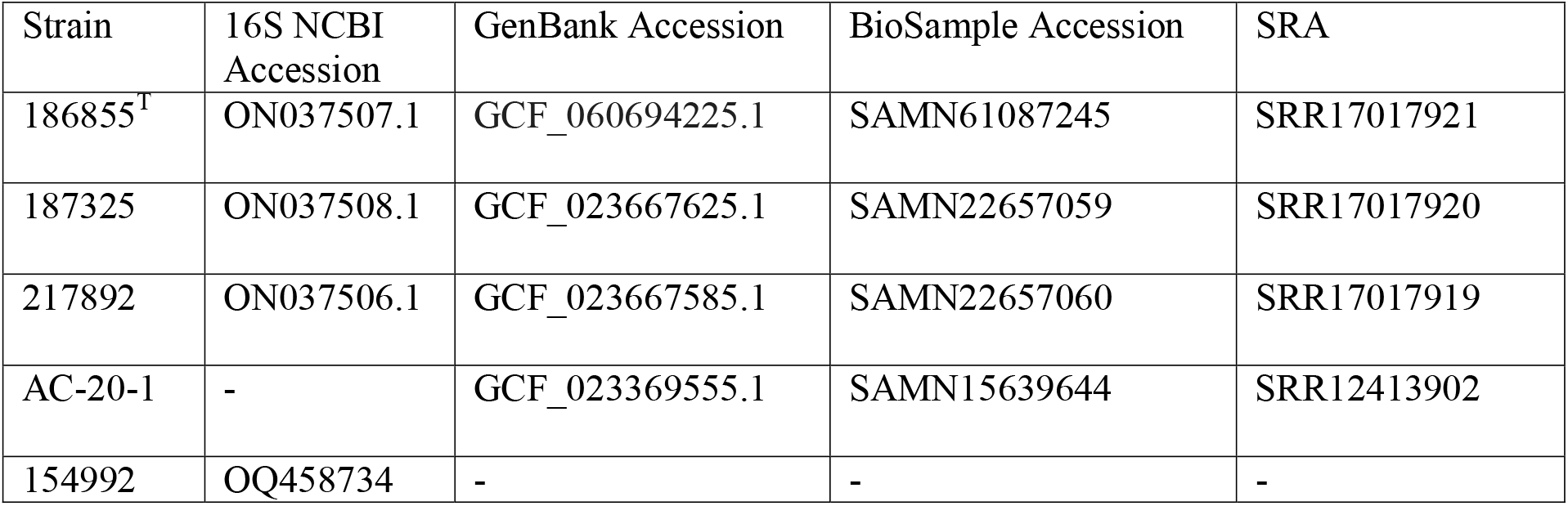

### DESCRIPTION OF *ACTINOMYCES CRICETOMYIS* SP. NOV

*Actinomyces cricetomyis* (cri.ce.to’my.is. N.L. gen. n. *cricetomyis*, of the African giant pouched rat *Cricetomys*). Bacterial cells are Gram-stain positive, rod-shaped and branch-forming bacilli, non-spore forming, non-motile and facultatively anaerobic. Colonies are white, raised and non-hemolytic when grown under aerobic conditions, observed as a haze of tiny growth at 24 hours and forming distinct colonies by 48 hours of incubation. It grows at 30°C - 42°C aerobically on trypticase soy agar with 5% sheep blood; at 35°C under anaerobic conditions on anaerobic *Brucella* blood agar; and at 42°C on *Campylobacter* selective agar under microaerophilic conditions. The optimal growth conditions for this organism are at 35°C with 6% CO_2_ on trypticase soy agar with 5% sheep blood, or at 35°C on *Campylobacter* selective agar under microaerophilic conditions. When assayed in the API ZYM systems, cells are positive for leucine arylamidase, α-galactosidase, β-galactosidase, α-glucosidase, β-glucosidase, and N-acetyl β-glucosaminidase however negative reactions for alkaline phosphatase, esterase lipase (C 8), lipase (C 14), valine arylamidase, cystine arylamidase, trypsin, α-chymotrypsin, acid phosphatase, naphthol-AS-BI-phosphohydrolase, α-mannosidase, and α-fucosidase. In contrast to closely related *Actinomyces* species, *A. cricetomyis* cells are weakly positive for β-glucuronidase. The Rapid ID 32A test show positive results for α-galactosidase, β-galactosidase, α-glucosidase, β-glucosidase, α-arabinosidase, arginine arylamidase, proline arylamidase, leucyl glycine arylamidase, phenylalanine arylamidase, leucine arylamidase, alanine arylamidase, glycine arylamidase, and serine arylamidase. In the Rapid ID 32A test, cells are heterogenous for mannose fermentation, raffinose fermentation, and histidine arylamidase. Negative results are reported for arginine dihydrolase, β-galactosidase 6-phosphate, β-glucuronidase, N-acetyl β-glucosaminidase, glutamic acid decarboxylase, α-fucosidase, nitrate reduction, indole production, alkaline phosphatase, pyroglutamic acid arylamidase, and glutamyl glutamic acid arylamidase in the Rapid ID 32A test. Notably, all *A. cricetomyis* strains tested are negative for tyrosine arylamidase in contrast to positive results for other related *Actinomyces* species. Cells are positive for catalase, aesculin hydrolysis, and acid production in phenol red (PR) dextrose and xylose. Cells are negative for urease, gelatin hydrolysis, and acid production in PR arabinose, mannitol, and trehalose. The genome size of the type strain is 2.7 Mb and the DNA G+C content is 72.6 mol%.

The type strain 186855^T^ (ATCC TSD-310^T^=LMG 32803^T^) was isolated from the ear of African giant pouched rats (*Cricetomys ansorgei*). The 16S rRNA gene accession of the type strain is ON037507 and the genome accession is GCF_060694225.1. The genome of the type strain can also be found separately at https://genomes.atcc.org/genomes/c202f05248c5406e.

## Acknowledgments

The authors wish to thank Profs. Aharon Oren and Bernhard Schink for consultation on the Latin spelling of the proposed species.

## Ethical approval

All work and maintenance of the animal colony followed American Society of Mammologists guidelines [22] and was approved under US Army Medical Research and Materiel Command (USAMRMC) Animal Care and Use Review Office (ACURO) and the Cornell University Institutional Animal Care and Use Committee. The samples used for this study were all collected as part of routine medical care of the animals.

